# Ectopic HOTTIP Expression Induces Non-canonical Transactivation Pathways to Promote Growth and Invasiveness in Pancreatic Ductal Adenocarcinoma

**DOI:** 10.1101/812800

**Authors:** Chi Hin Wong, Chi Han Li, Qifang He, Stephen Lam Chan, Joanna Hung-Man Tong, Ka-Fai To, Yangchao Chen

**Author notes:** These authors contributed equally (Chi Hin Wong, Chi Han Li). **Contact Information** Yangchao Chen, PhD, School of Biomedical Sciences, Faculty of Medicine, Chinese University of Hong Kong, Shatin, Hong Kong.

## Abstract

LncRNA HOTTIP (HOXA transcript at the distal tip) is upregulated in pancreatic ductal adenocarcinoma (PDAC). However, oncogenic pathway mediated by HOTTIP is not fully understood. We identified canonical HOTTIP-HOXA13 targets: CYP26B1, CLIC5, CHI3L1 and UCP2 which were responsible for cell growth and cell invasion. Importantly, genome-wide analysis revealed that 38% of the genes regulated by HOTTIP contained H3K4me3 and HOTTIP enrichment at their promoters, without HOXA13 binding. HOTTIP complexed with WDR5-MLL1 to *trans*-activate oncogenic proteins CYB5R2, SULT1A1, KIF26A, SLC1A4 and TSC22D1 through directly triggering H3K4me3 at their promoters. The WDR5, MLL1 and H3K4me3 levels at their promoters and their expression levels were sensitive to HOTTIP overexpression and knockdown. These suggested the importance of non-canonical *trans*-acting HOTTIP-WDR5-MLL1 pathway in HOTTIP-regulatory mechanism by promoting the expression of oncogenic proteins. Furthermore, we dissected the mechanism by which HOTTIP was regulated by miR-497 in PDAC. Analysis of PDAC human tissues also revealed that HOTTIP was negatively correlated with miR-497 level. Our findings demonstrated that HOTTIP, upregulated in PDAC due to the loss of inhibitory miR-497, promoted PDAC progression through canonical HOTTIP-HOXA13 axis and a novel non-canonical *trans*-acting HOTTIP-WDR5-MLL1-H3K4me3 pathway.

## 1. Introduction

Pancreatic ductal adenocarcinoma (PDAC) is aggressive and is the fourth leading cause of cancer death [1]. It is predicted that both the incidence and the mortality keep increasing in the following two decades [2]. Due to the lack of reliable diagnostics and therapeutics, most patients die within a year after diagnosis. Even worse, the current therapies can only extent the lifespan by few months. Therefore, it is of urgent significance to understand the mechanism of PDAC tumorigenesis to develop effective therapies. Studies found that most of the genome encodes RNAs with no protein-coding ability, termed as non-coding RNAs. Among the diverse types of non-coding RNAs, the functions of microRNAs (miRNAs), with the size less than 200 bp, are extensively characterized. Another group of non-coding RNAs with more than 200 bp, termed long non-coding RNAs (lncRNAs), is now getting more attentions. lncRNAs play important roles in many cellular processes including chromatin modifications and alternative splicing [3,4]. HOXA transcript at the distal tip (HOTTIP), which locates at the 5’ tip of HOXA gene clusters, binds to the adaptor protein WDR5 and H3K4 methyltransferase MLL1 [5]. Chromosomal looping brings this HOTTIP-WDR5-MLL1 complex close to 5’ HOXA genes especially HOXA13 and promotes their expressions [6]. The HOTTIP-HOXA13 axis plays an important role in cancer, such as colorectal cancer, gastric cancer, and hepatocellular carcinoma [7–9]. HOTTIP-HOXA13 axis also promoted PDAC development and gemcitabine resistance [10].

In this study, we performed genome-wide profiling of lncRNAs expression in PDAC tumors by microarray. We found that HOTTIP was significantly upregulated in PDAC cell lines and primary tumors and promoted tumor growth and invasion. We identified the effectors of HOTTIP-HOXA13 axis: Cytochrome P450 26B1 (CYP26B1), Chloride intracellular channel 5 (CLIC5), Chitinase 3-like 1 (CHI3L1) and Uncoupling Protein 2 (UCP2), which participated in PDAC progression. Importantly, we are the first to identify the non-canonical *trans*-acting WDR-MLL1-HOTTIP pathway, which HOTTIP directly activated genes in different chromosomes including Cytochrome B5 Reductase 2 (CYB5R2), Sulfotransferase Family 1A Member 1 (SULT1A1), Kinesin Family Member 26A (KIF26A), Solute Carrier Family 1 Member 4 (SLC1A4) and TSC22 Domain Family Member 1 (TSC22D1), without the involvement of HOXA13. Also, we identified miR-497 as the regulator of HOTTIP in PDAC.

## 2. Methods and materials

### 2.1 Clinical Sample and Cell lines

Sixty pairs of PDAC primary tumor and adjacent non-tumor tissues were obtained from patients who underwent pancreatic resection at the Prince of Wales Hospital, Hong Kong. All specimens were fixed and embedded into paraffin. The study was carried out with the approval of the Joint CUHK-NTEC Clinical Research Ethics Committee. PDAC Cell lines PANC-1, SW1990, CAPAN-2, CFPAC-1, PANC0403, and BxPC-3 were obtained from American Type Culture Collection. Human pancreatic ductal epithelial (HPDE) cell line was a gifted from Dr Ming-Sound Tsao (University Health Network, Ontario Cancer Institute and Princess Margaret Hospital Site, Toronto) [11]. All cell lines were cultured under the condition as previously described [12].

### 2.2 Microarray analysis

Microarray analysis of lncRNA expression on 4 pairs of PDAC tumor samples was performed on Arraystar Human LncRNA Microarray V4.0 platform. Microarray data are available at ArrayExpress E-MTAB-7305. To identify genes which are regulated by HOTTIP, microarray analysis after knockdown of HOTTIP in SW1990 cells was performed on Affymetrix GeneChip Human Genome U133 Plus 2.0 Array platform.

### 2.3 Plasmid and oligonucleotide transfection

Plasmid overexpressing HOTTIP was purchased from Applied Biological Materials Inc (British Columbia) and was transfected into HPDE cells using lipofectamine 3000 (Invitrogen) according to the manufacturer’s protocol. The siRNAs targeting HOTTIP, HOXA13, WDR5, MLL1, MLL2, MLL3, CYP26B1, CYB5R2, UCP2, SULT1A1, CLIC5, and CHI3L1; miRNA mimics and miRNA inhibitors were purchased from GenePharma. The sequences of siRNAs and miRNAs used in this study are shown in Supplementary Table S2. The oligonucleotides were transfected into the PDAC cells using Lipofectamine 3000. After 72 hrs, cells were collected for the following experiments. The pmiR-HOTTIP reporter was constructed by cloning HOTTIP coding sequence with predicted binding sites for miR-497 into pmiR-REPORTER luciferase plasmid (Ambion).

### 2.4 Western blot analysis

Whole cell extract was prepared by lysing cells in NP-40 lysis buffer with proteinase inhibitors (Roche) and phosphatase inhibitor (Thermofisher). Protein concentration was determined by BCA assay (Pierce). Proteins were separated by SDS-PAGE, transferred to PVDF membrane and immunoblotted overnight at 4 °C with 1:1000 primary antibodies against HOXA13 (ab106503; abcam), MLL1 (#ABE240; Millipore), MLL2 (#ABE206, Millipore), MLL4 (NB600-254, Novus Biologicals), WDR5 (ab56919; abcam), CYB5R2 (H00051700-B02P; abnova), CYP26B1 (H00056603-M01; abnova), UCP2 (ab97931; abcam) and GAPDH (#5174; Cell signaling). Then, the blots were washed three times with TBST, followed by incubation with 1:2000 secondary anti-rabbit antibody or secondary anti-mouse antibody at room temperature for 1 hr.

### 2.5 Luciferase assay

pmiR-HOTTIP reporter or pmiR-REPORTER was co-transfected with miR-497 or miR-NC mimics into HEK293 cells. The luciferase plasmid pNL1.1 TK was used as an internal control. After 72hrs, the luciferase activity was measured using Nano-Glo Dual luciferase reporter system (Promega) with SpectraMax® i3x Multi-Mode Microplate Detection Platform. The ratio of Firefly luciferase to NanoLuc luciferase was calculated for each sample and was normalized using pmiR-REPORTER control.

### 2.6 Additional methods

Quantitative reverse-transcription PCR (qRT-PCR), immunohistochemistry staining, MTT cell viability assay, invasion assay, Chromatin Immunoprecipitation (ChIP) assay, and chromatin Isolation by RNA Purification (ChIRP) assay, are described in Supplementary Materials and Methods.

### 2.7 Statistical analysis

GrapPad Prism 6 (GraphPad Software Inc., La Jolla, CA) was used for statistical analysis. All data were presented as mean ± SD. Student’s t-test was used to analyze the differences between two groups. Pearson’s correlation coefficient was used to analyze the correlation between groups. P value less than 0.05 was considered statistically significant.

## 3 Results

### 3.1 HOTTIP promotes PDAC proliferation and invasion

To investigate the roles of differentially expressed lncRNAs in progression and development of PDAC, we profiled their expressions using microarray (Human LncRNA microarray V2.0), which interrogated 33,045 lncRNAs and 30,215 protein-coding genes simultaneously, on 4 pairs of PDAC primary tumor and paired non-tumor tissues. Microarray analysis indicated that HOTTIP was upregulated by at least 2-fold in all 4 patients (Supplementary Figure S1A). Quantitative real-time polymerase chain reaction (qRT-PCR) in 60 paired PDAC primary tumors confirmed that HOTTIP expression level was frequently higher in PDAC, compared to adjacent non-tumor tissues (Supplementary Figure S1B). Also, analysis of The Cancer Genome Atlas (TCGA) dataset also revealed up-regulation of HOTTIP in PDAC (Supplementary Figure S1C). In addition, we measured the HOTTIP level in a panel of PDAC cells including PANC-1, SW1990, Capan-2, CFPAC-1, PANC0403 and BxPC-3. Compared to non-tumorigenic human pancreatic ductal epithelial (HPDE) cells, HOTTIP was uniformly and significantly overexpressed in PDAC cell lines (Supplementary Figure S1D).

To characterize the role of HOTTIP in PDAC, we first utilized knockdown approach to determine whether treating PDAC cells with siRNA against HOTTIP could inhibit cancer development (Supplementary Figure S1E). Indeed, knockdown of HOTTIP inhibited cell growth, as assessed by MTT assay (Supplementary Figure S1F). In addition, transwell cell invasion assay revealed that knockdown of HOTTIP hindered PDAC cell invasion (Supplementary Figure S1G). Next, we investigated the oncogenic potential of HOTTIP through overexpression approach in HPDE cells (Supplementary Figure S1H). Overexpression of HOTTIP promoted cell growth in HPDE cells (Supplementary Figure S1I). Collectively, these findings suggested that the upregulation of HOTTIP in PDAC promoted cancer progression.

### 3.2 Identification of HOTTIP targets in PDAC

To unveil the molecular mechanism by which HOTTIP promoted PDAC growth and invasion, we performed microarray (GeneChip Human Genome U133 Plus 2.0 Array, Affymetrix) to identify deregulated genes after HOTTIP knockdown in SW1990 cells. Gene Ontology (GO) and Kyoto Encyclopedia of Genes and Genomes (KEGG) pathway analysis revealed an enrichment of genes participating in transcription, cell fate determination, protein binding and metabolism (Supplementary Table S3 and S4). It has been reported that HOTTIP promoted gene expression through the induction of activating histone markers H3K4 methylation, and the targets of HOTTIP were transcriptional activator HOXA family members [5], therefore we focused on genes that were down-regulated upon inhibition of HOTTIP. We found that the expression levels of CHI3L1, CLIC5, CYB5R2, CYP26B1, SULT1A1 and UCP2 were significantly decreased after HOTTIP knockdown (Fig. 1A and Supplementary Figure S2), while they were upregulated when HOTTIP was overexpressed in HPDE cells (Fig. 1B). Then we analyzed their expression levels in PDAC cells and primary tumors. As expected, CHI3L1, CLIC5, CYB5R2, CYP26B1, SULT1A1 and UCP2 were significantly upregulated in both PDAC cells and primary tumors (Fig. 1C and Supplementary Figure S3A). In addition, there was a positive correlation between the mRNA or protein levels of these downstream effectors and HOTTIP expression level in human PDAC primary tumors (Fig. 1D and E).

**Fig. 1.**
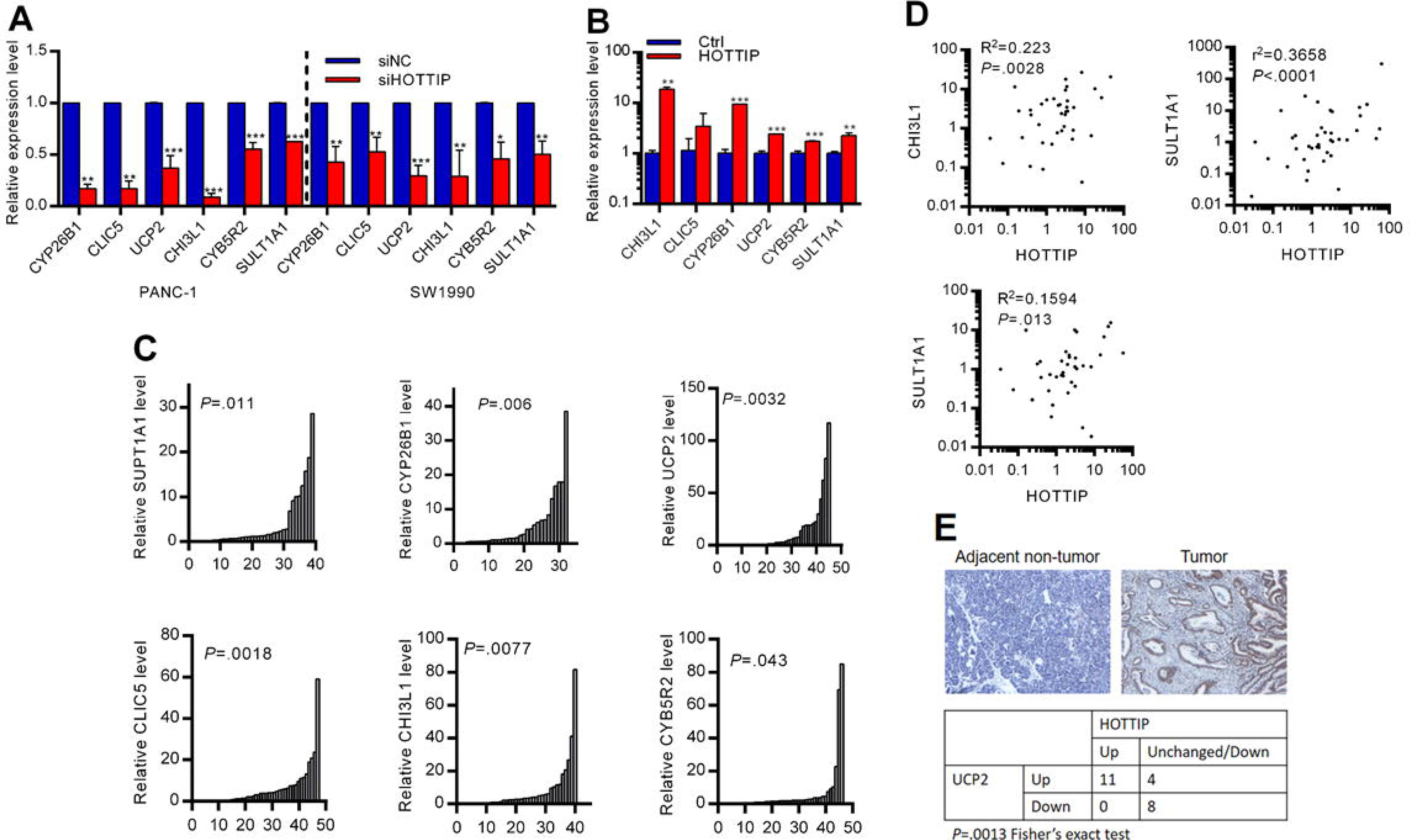
Identification of HOTTIP target genes. **A-B**, Expression of HOTTIP target genes (**A**) was inhibited in PDAC cells treated with siHOTTIP; and (**B**) was upregulated in non-tumorigenic human pancreatic ductal epithelial (HPDE) cells when HOTTIP was overexpressed (n=3). **C**, Expression levels of HOTTIP targets in PDAC FFPE samples, compared to adjacent non-tumor tissues (n=37-47). **D**, The correlation between CHI3L1, SULT1A1, CLIC5 and HOTTIP expression in PDAC tumor samples. **E**, Representative images of UCP2 immunohistochemistry on PDAC samples (right panel), compared with adjacent non-tumor tissue (left panel). UCP2 protein level was correlated with HOTTIP expression (lower panel). *P < 0.05; **P < 0.01; ***P < 0.001 compared with control.

Although these HOTTIP target genes were reported to play important roles in cancer, their functions in PDAC are not fully understood. Therefore, we utilized the knockdown approach by siRNAs to explore their functions in PDAC (Supplementary Figure S3B and C). Knockdown of UCP2, CYB5R2, CLIC5 and CYP26B1 resulted in a significant reduction in PDAC cell growth (Fig. 2A-D). Moreover, depletion of CYB5R2, CHI3L1, SULT1A1 and CLIC5 inhibited cell invasion (Fig. 2E). Collectively, our results suggested that CHI3L1, CLIC5, CYB5R2, CYP26B1, SULT1A1 and UCP2 played important roles in PDAC progression.

**Fig. 2.**
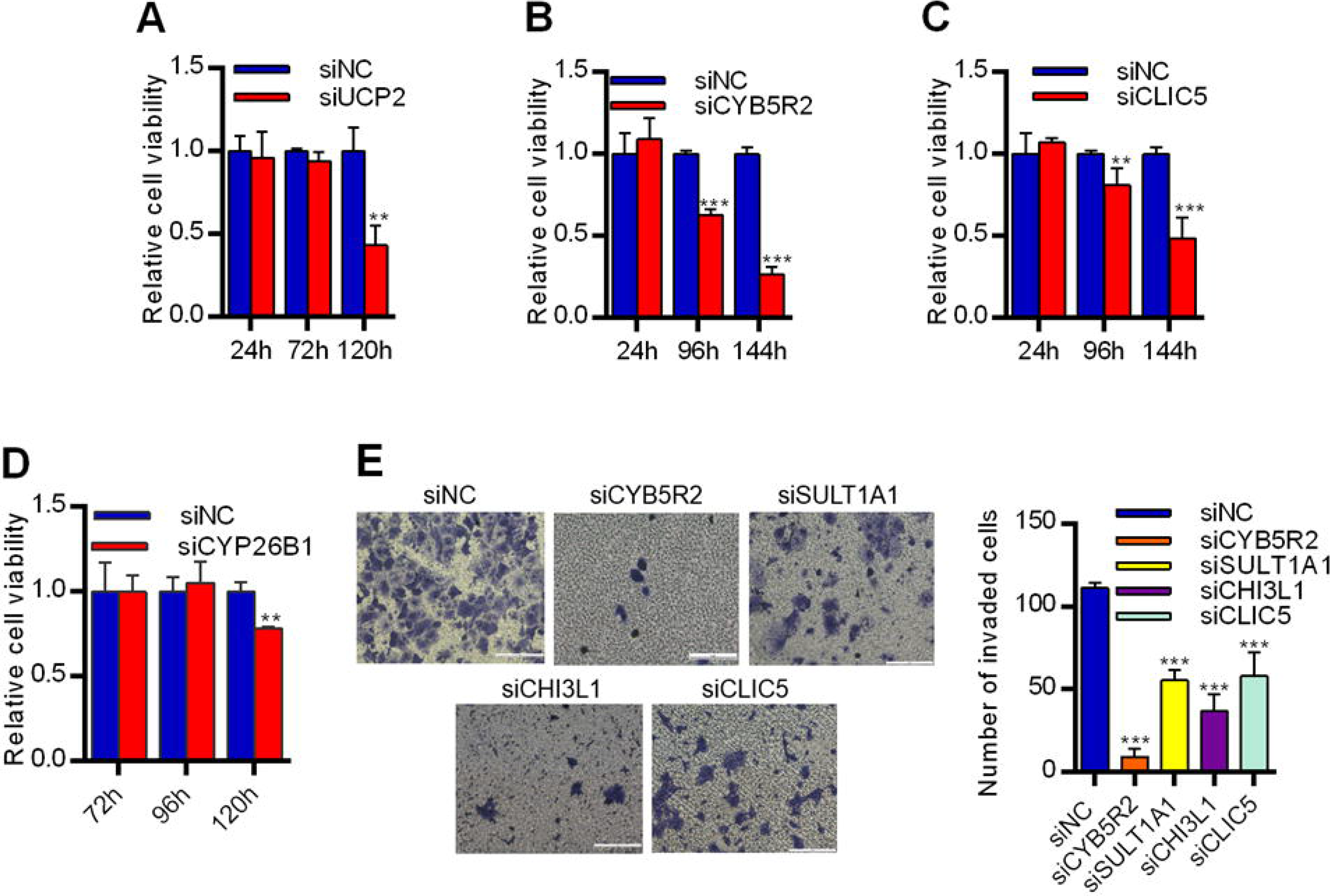
HOTTIP target genes promote cell growth and invasion in PDAC. **A-D**, Inhibition of cell viability in PANC-1 cells treated with (**A**) siUCP2, (**B**) siCYB5R2, (**C**) siCLIC5 and (**D**) siCYP26B1 (n=4). **E**, Knockdown of CHI3L1, CLIC5, CYB5R2 and SULT1A1 inhibited cell invasiveness in PANC-1 cells. *P < 0.05; **P < 0.01; ***P < 0.001 compared with control.

### 3.3 Canonical HOTTIP-HOXA13 pathway in PDAC

Next, we further investigated the underlying mechanism. HOTTIP activated several 5’ HOXA genes in human primary fibroblast during development [5]. Therefore, we measured the expression level of HOXA genes cluster after HOTTIP depletion. Contrasting to previous finding, HOXA13 was down-regulated after knockdown of HOTTIP (Supplementary Figure S4A). HOTTIP also bound the promoter of HOXA13 gene, demonstrated by ChIRP assay (Supplementary Figure S4B). Overexpression of HOTTIP increased the occupancies of MLL1, WDR5 and H3K4me3 on HOXA13 promoter (Supplementary Figure S4C). On the other hand, knockdown of HOTTIP partner MLL1 reduced its occupancy to the promoter, and in turn hindered the expression of HOXA13 gene (Supplementary Figure S4D and E). We then measured HOXA13 expression level in a panel of PDAC cell lines. We found the HOXA13 was significantly upregulated in PDAC cells (Supplementary Figure S4F). Analysis of TCGA datasets also revealed that HOXA13 was upregulated and was positively correlated with HOTTIP expression in PDAC (Supplementary Figure S4G). Also, knockdown of HOXA13 by siRNA significantly inhibited the PDAC cell growth (Supplementary Figure S4H and I). These results demonstrated the importance of HOTTIP-HOXA13 pathway in PDAC.

To identify the downstream targets under HOTTIP-HOXA13 axis, we knocked down HOXA13. We found that CHI3L1, CLIC5, CYP26B1, and UCP2 were down-regulated after knockdown of HOXA13 (Fig. 3A and B). Interestingly, CYB5R2 and SULT1A1 were not sensitive to HOXA13 knockdown. Since HOXA13 is DNA-binding transcription factor, we hypothesized that HOXA13 mediated the expression of CHI3L1, CLIC5, CYP26B1, and UCP2 through binding to their promoters. As expected, we observed the enriched occupancy of HOXA13 on the promoter region of these down-stream targets in SW1990 cells (Fig. 3C). Knockdown of HOTTIP or HOXA13 significantly reduced the occupancy of HOXA13 on the promoter of CHI3L1, CLIC5, CYP26B1, and UCP2 (Fig. 3D and E). Similarly, overexpression of HOTTIP in HPDE cells promoted the binding of HOXA13 on the promoters of CHI3L1, CLIC5, CYP26B1, and UCP2 (Fig. 3F). Altogether, these results demonstrated that CHI3L1, CLIC5, CYP26B1, and UCP2 were regulated under canonical HOTTIP-HOXA13 axis.

**Fig. 3.**
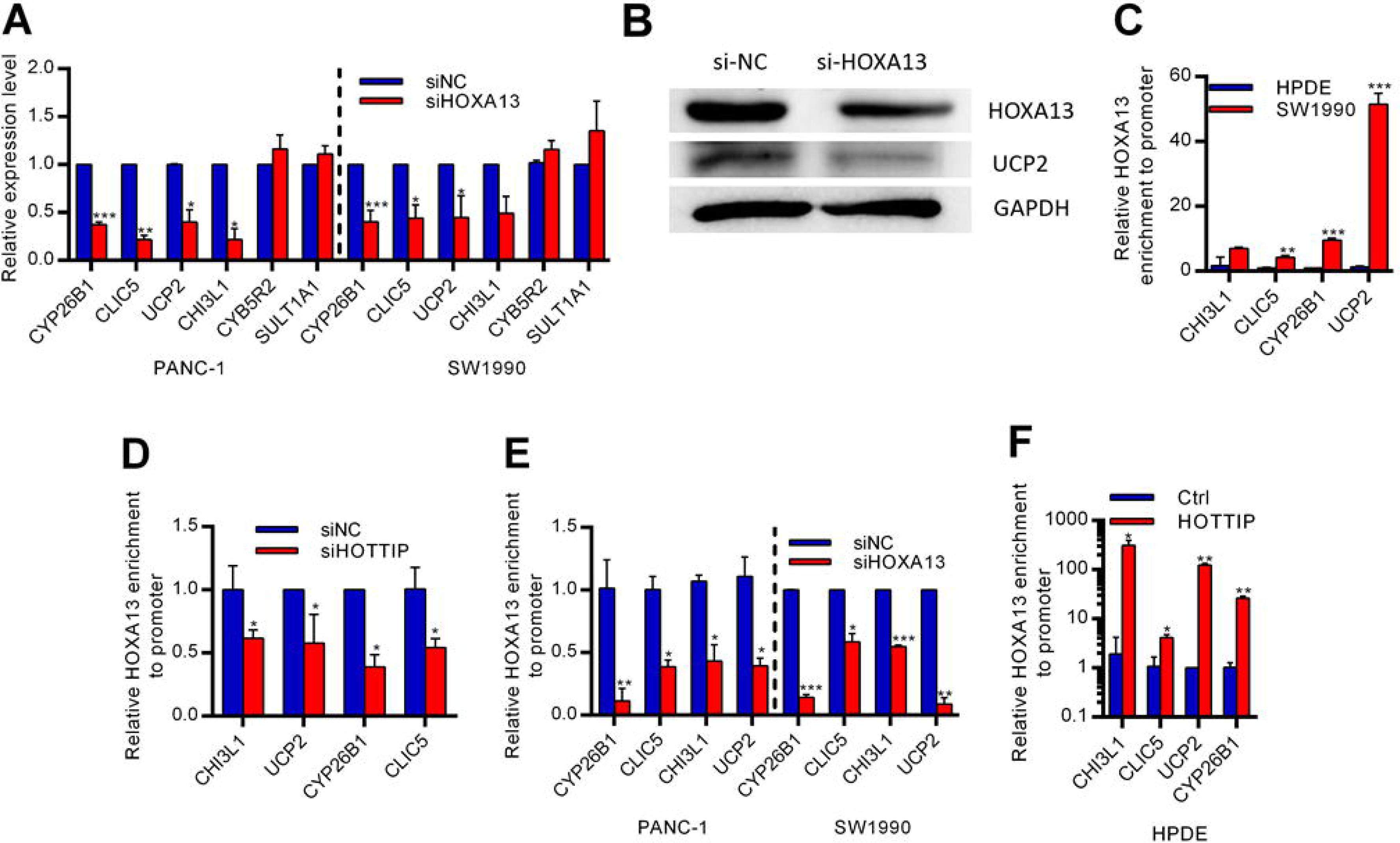
Identification of HOTTIP target genes regulated by canonical HOTTIP-HOXA13 axis. **A**, Expression levels of HOTTIP target genes in PDAC cells treated with siHOXA13 (n=3). **B**, Western blot analysis of HOTTIP targets in PANC-1 cells treated with siHOXA13. **C**, ChIP analysis of HOXA13 binding to the promoter of HOTTIP target genes (n=3). **D-F**, HOXA13 occupancy to the promoter of CHI3L1, CLIC5, CYP26B1 and UCP2 after knockdown of (**D**) HOTTIP and (**E**) HOXA13 in PDAC cells (n=3); and (**F**) after overexpression of HOTTIP in HPDE cells (n=3). *P < 0.05; **P < 0.01; ***P < 0.001 compared with control.

### 3.4 Genome-wide identification of non-canonical HOTTIP-WDR5-MLL1 pathway

Since CYB5R2 and SULT1A1 were not sensitive to HOXA13 knockdown, they may be regulated by HOXA13-independent pathway. Since HOTTIP activates HOXA13 through WDR5-MLL1 mediated H3K4me3, we hypothesized that HOTTIP downstream targets were regulated by the non-canonical WDR5-MLL1-HOTTIP pathway. Genome-wide analysis was performed using publicly available H3K4me3 (GSE85886) and HOXA13 (GSE49402) dataset and HOTTIP binding analysis by Triplex Domain Finder to identify genes with HOTTIP and H3K4me3 but not HOXA13 enrichment at their promoters (Fig. 4A) [14–16]. As expected, HOTTIP targets CYB5R2, SULT1A1, KIF26A, SLC1A4 and TSC22D1 were insensitive to HOXA13 knockdown (Fig. 3A and 4B). Also, there was a decrease in expression levels of CYB5R2 and SULT1A1 after knockdown of WDR5 or MLL1 (Fig. 4C-E). The HOTTIP-induced CYB5R2 and SULT1A1 expression can be rescued by WDR5 or MLL1 knockdown (Supplementary Figure SS5A). Also, we observed increased occupancies of HOTTIP, MLL1, WDR5 and H3K4me3 on the promoter of CYB5R2 and SULT1A1 in PDAC cells, compared to HPDE cells (Fig. 4F and G). Knockdown of HOTTIP, MLL1 or WDR5 significantly reduced their occupancies and H3K4me3 level on the promoter of CYB5R2 and SULT1A1 (Supplementary Figure SS5B-D). Similarly, overexpression of HOTTIP in HPDE cells increased the occupancy of MLL1 and WDR5, and H3K4me3 level on the promoters of CYB5R2, KIF26A, SLC1A4 and TSC22D1 (Fig. 5H). These results suggested that CYB5R2, KIF26A, SLC1A4, TSC22D1 and SULT1A1 were regulated under the physical interaction of HOTTIP, MLL1 and WDR5 on their promoters. Although HOTTIP also interacted with other MLL family members such as MLL2, knockdown of MLL2, MLL3 or MLL4 did not consistently inhibited the expression of CYB5R2 and SULT1A1 in PDAC cells (Supplementary Figure S6). Collectively, these data implied that CYB5R2, SULT1A1, KIF26A, SLC1A4 and TSC22D1 were regulated by H3K4me3 on their promoters under non-canonical HOTTIP-WDR5-MLL1 complex.

**Fig. 4.**
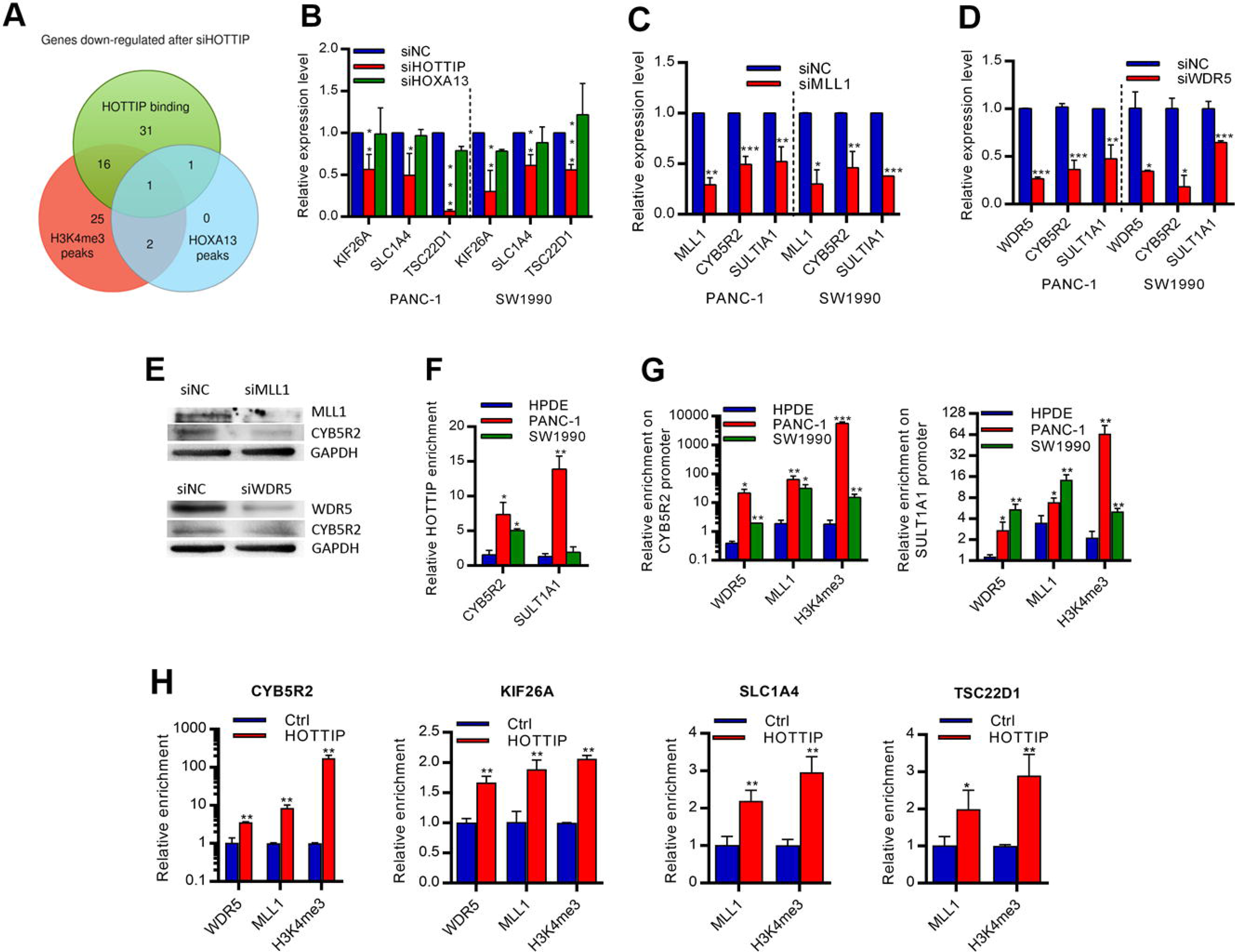
Identification of HOTTIP target genes regulated by non-canonical HOTTIP-WDR5-MLL1 pathway. **A**, Genome-wide analysis of H3K4me3, HOXA13 and HOTTIP enrichment to the promoters of HOTTIP-regulated genes. **B**, Identification of HOTTIP downstream targets which were not sensitive to HOXA13 knockdown (n=3). **C**, Expression levels of HOTTIP target genes in PDAC cells after the knockdown of H3K4 methyltransferase MLL1 (n=3). **D**, CYB5R2 and SULT1A1 expressions were reduced in PDAC cells treated with siWDR5 (n=3). **E**, Western blot analysis of CYB5R2 after the knockdown of MLL1 and WDR5 in PANC-1 cells. **F**, ChIRP assay analysis of HOTTIP binding to the promoter of CYB5R2 and SULT1A1 in PANC-1 and SW1990 cells. Data were normalized to negative control and were compared to HOTTIP enrichment levels in HPDE cells (n=3). **G**, ChIP assay analysis revealed the occupancies of WDR5, MLL1 and H3K4me3 at the promoter of CYB5R2 and SULT1A1 in PDAC cells. Data were compared to HOTTIP enrichment levels in HPDE cells (n=3). **H**, WDR5, MLL1 and H3K4me3 levels at the promoter of CYB5R2, KIF26A, SLC1A4 and TSC22D1 were increased when HOTTIP was overexpressed in HPDE cells (n=3). *P < 0.05; **P < 0.01; ***P < 0.001 when compared with control.

**Fig. 5.**
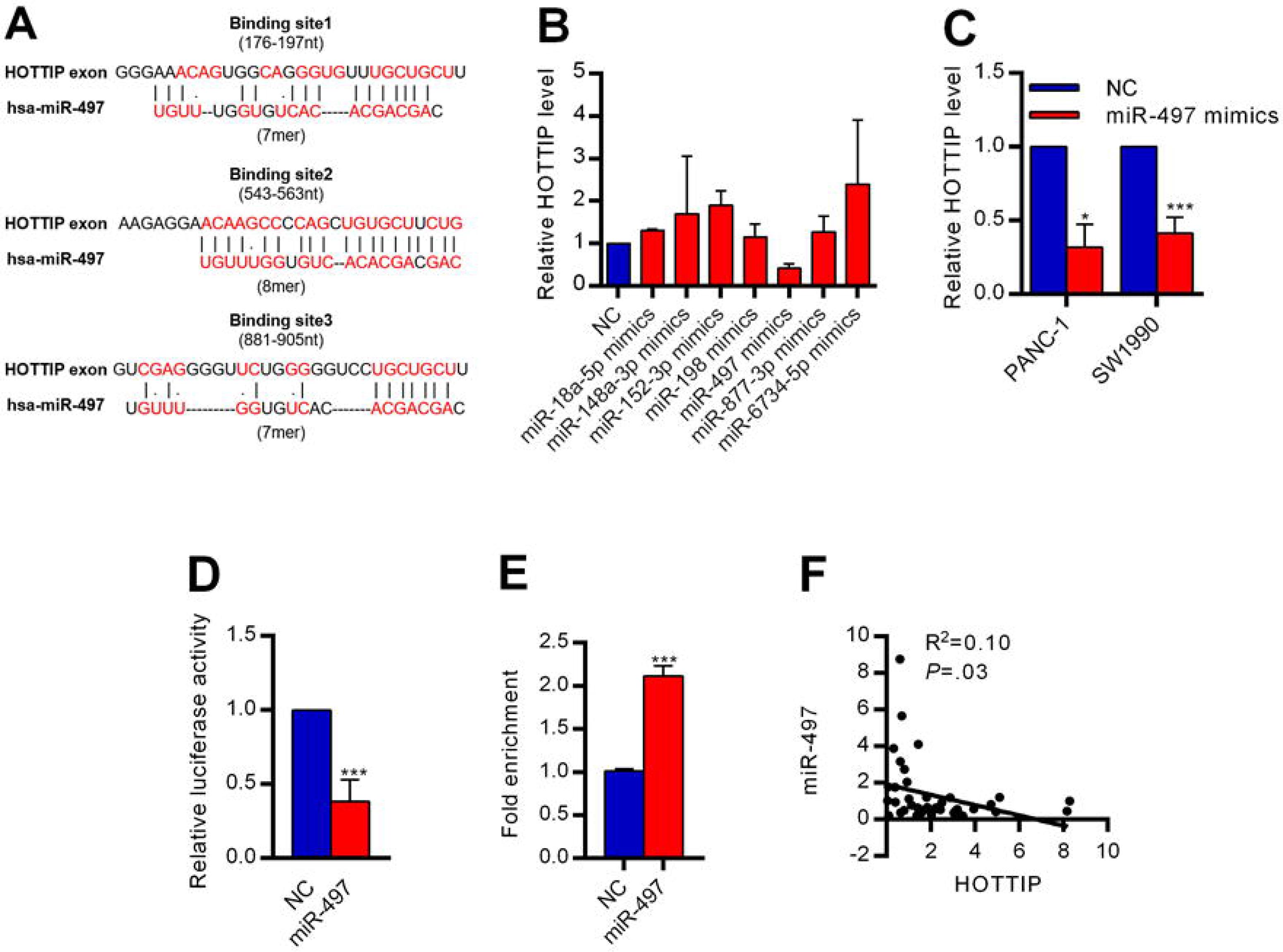
miR-497 regulates HOTTIP in PDAC cells. **A**, Bioinformatics analysis was used to identify potential HOTTIP regulator, based on the predicted binding sequences on the coding region of HOTTIP. The predicted binding sites of miR-497 were shown. **B**, Validation experiment was performed by transfecting miRNA mimics in SW1990 cells (n=2). **C**, PDAC cells were transfected with miR-497 mimics and HOTTIP expression was measured using real-time qPCR (n=3). **D**, miR-497 interacted with the coding region in HOTTIP, demonstrated by luciferase assay (n=3). **E**, miRNA pull down assay revealed miR-497 interacts with HOTTIP (n=3). **F**, The correlation between HOTTIP and miR-497 (n=43) expression in human PDAC tissue samples. *P < 0.05; **P < 0.01; ***P < 0.001 when compared with control.

### 3.5 miR-497 regulates HOTTIP expression in PDAC

To determine the mechanism by which miRNA regulate HOTTIP expression, we first searched the HOTTIP exon regions for putative binding sites of cancer-associated miRNAs using bioinformatics prediction tool (Fig. 5A) [17]. Then, the cancer-associated miRNAs that regulate HOTTIP in PDAC were validated by transfecting each miRNA mimics in PDAC cells. miR-497 had significantly reduced HOTTIP expression (Fig. 5B and C). Luciferase assay and miRNA pull down assay also revealed the binding of miR-497 to the HOTTIP exon regions (Fig. 5D and E). These results indicated that miR-497 directly modulated HOTTIP expression through interaction with the exon regions, but not 5’ and 3’ intronic regions. To study the significance of miR-497 on HOTTIP expression in PDAC, we examined miR-497 expression in PDAC primary tumors. miR-497 was significantly inversely correlated with HOTTIP expression (Fig. 5F). Collectively, these results indicated that miR-497 could directly bind to HOTTIP exon region and regulate HOTTIP expression in PDAC.

## 4 Discussion

Overexpression of HOTTIP promotes the development of cancer including pancreatic cancer [10,18]. Gene Ontology (GO) and Kyoto Encyclopedia of Genes and Genomes (KEGG) pathway analysis revealed an enrichment of genes participating in transcription, cell fate determination, protein binding and metabolism after knockdown of HOTTIP. These processes and pathways are strongly associated with cancer [19], suggesting that HOTTIP plays critical roles in PDAC. HOTTIP induces the expression of HOXA13 through WDR5-MLL1-mediated H3K4me3 in the promoter of HOXA13 gene [5]. Studies suggested that HOTTIP-HOXA13 axis mediates tumorigenesis in many types of cancer such as hepatocellular carcinoma and lung cancer [9, 20]. Also, a recent study demonstrated that in additional to epigenetic regulation of HOXA13, HOTTIP promotes HOXA13 expression through eliminating HOXA13-inhibiting miR-30b in esophageal squamous carcinoma [21]. However, the HOTTIP-HOXA13 regulatory pathway is not fully understood in pancreatic cancer. There is a positive association between HOTTIP and HOXA13 expression in PDAC tumors. Also, knockdown of HOTTIP suppresses the expression of HOXA13 [10]. However, it has also been reported that inhibiting HOTTIP does not affect the expression of HOXA13, but down-regulates other HOXA genes such as HOXA10 [18]. Clearly, more studies are needed to reveal the molecular mechanism of HOTTIP in regulating the HOXA genes during PDAC carcinogenesis. In this study, we found that HOXA13 was regulated by HOTTIP in PDAC. Importantly, we revealed that HOTTIP bind to HOXA13 promoter and mediated gene-activating H3K4me3 through WDR5-MLL1 complex.

Notably, we here discovered the downstream target genes of HOTTIP-HOXA13 axis; CHI3L1, CYP26B1, CLIC5 and UCP2. HOXA13 promoted the expression of its targets via binding to their promoters. Studies showed that CHI3L1, CYP26B1, CLIC5 and UCP2 are associated with the development of a panel of cancers. CHI3L1, a secreted glycoprotein, plays an important role in carcinogenesis. Upregulation of CHI3L1 promoted cell proliferation, invasion and drug resistance in cancer, including colorectal cancer, glioma, and papillary thyroid carcinoma [22–24]. CLIC5 promoted cell proliferation in prostate cancer and functioned as metastatic marker in breast cancer [25]. CLIC family member CLIC1 was also expressed in PDAC tumors and prostate cancer, and promoted cancer proliferation [26,27]. UCP2 protected colorectal cancer cells from apoptosis [28]. However, UCP2 functioned as tumor suppressor in glioma through mediating mitochondrial retrograde signaling [29]. Their roles in PDAC development are not fully understood. In this study, we found that CHI3L1, CYP26B1, CLIC5 and UCP2 were upregulated in PDAC cells and primary tumors, and were positively correlated to HOTTIP expression. Importantly, CHI3L1, CYP26B1, CLIC5 and UCP2 promoted cell growth and invasion. Therefore, we demonstrated that HOTTIP mediated the expression of HOXA13, which in turn promoted PDAC cell growth through promoting the expression of CHI3L1, CYP26B1, CLIC5 and UCP2 via binding to their promoters.

The *cis*-acting HOTTIP on HOXA genes was comprehensively illustrated in many studies. Meanwhile, our research is the first to reveal the novel *trans*-acting roles of HOTTIP in promoting PDAC development. Since CYB5R2 and SULT1A1 were not sensitive to HOXA13 knockdown, they may be the non-canonical targets of HOTTIP. HOTTIP formed complex with WDR5 and MLL1, resulting in activation of nearby HOXA genes through H3K4me3 [5,6]. Therefore, we hypothesized that CYB5R2 and SULT1A1 were under the regulation of HOTTIP-WDR5-MLL1 complex. Genome-wide analysis demonstrated that 38% of the genes regulated by HOTTIP in different chromosomes contained H3K4me3 and HOTTIP enrichment at their promoters. This suggested that the non-canonical *trans*-acting HOTTIP-WDR5-MLL1 pathway played an important role in HOTTIP regulatory pathway.

Our results also demonstrated that CYB5R2, SULT1A1, KIF26A, SLC1A4 and TSC22D1 were inhibited by knockdown of MLL1 and WDR5. HOTTIP, WDR5, MLL1, and H3K4me3 enrichment were also observed on the promoters of CYB5R2 and SULT1A1. Their occupancies on the promoters of CYB5R2, SULT1A1, KIF26A, SLC1A4 and TSC22D1 were sensitive to overexpression or depletion of HOTTIP, MLL1 and WDR5. Therefore, we here suggested that, in addition to HOXA genes, HOTTIP-WDR5-MLL1 complex also directly activated CYB5R2, SULT1A1, KIF26A, SLC1A4 and TSC22D1 through H3K4me3 on their promoters. It has been suggested that HOTTIP was brought close to HOXA genes through chromosome looping, and in turn mediated the expression of nearby HOXA7, HOXA9 and HOXA13. Therefore, in the past, many studies focused on HOTTIP-HOXA-mediated carcinogenesis [9, 10,17, 18, 30]. However, our findings demonstrated that HOTTIP could *trans*-activate multiples downstream targets, which were located at different chromosomes, through direct interaction to their promoters. HOTTIP was located at the chromosome 7, while CYB5R2, SULT1A1, KIF26A, SLC1A4 and TSC22D1 were located at chromosome 11, 16, 17, 2 and 13 respectively. This suggested that HOTTIP may regulate distant genes but not nearby HOXA gene clusters exclusively. It was possible that the regulating mechanism of HOTTIP in cancer cells was highly deviated from non-tumor counterparts, as there were about 102 to 104-fold increase of HOTTIP in cancer cells while it was reported that HOTTIP had a low copy number that enabled *cis*-regulation [5]. Therefore, chromosome looping may not be the sole mechanism for HOTTIP-mediated gene regulation. Since, HOTTIP is frequently upregulated in many cancers, activation of distal genes by direct HOTTIP binding to the promoters by the formation of RNA/DNA triplex may also be one of the major mechanisms in oncogenesis in other cancer types.

Since our findings demonstrated that HOTTIP played an important role in PDAC development; however, the regulation of HOTTIP is poorly understood. Therefore, we attempted to investigate the regulatory mechanism of HOTTIP. Previous studies demonstrated the miRNAs-mediated degradation of lncRNAs through perfect or partial base pairing [31–33,35–37]. miR-9 degraded lncRNA Metastasis Associated Lung Adenocarcinoma Transcript 1 (MALAT1) in nucleus, in Ago-2 dependent manner [31]. miR-21 suppressed the expression of lncRNA growth arrest-specific 5 (GAS5) [32]. Alternatively, miRNA could promote gene expression through direct base pairing or indirectly triggering promoter hypermethylation [33–35]. miR-140 stabilized lncRNA Nuclear Enriched Abundant Transcript 1 (NEAT1) by directly binding to exon region of NEAT1 [35]. Therefore, we hypothesized that the upregulation of HOTTIP might be contributed by the deregulated miRNA expression in PDAC. We found that miR-497, down-regulated in PDAC, was the post-transcriptional suppressor of HOTTIP. miR-497 was aberrantly inhibited in cancer, such as renal carcinoma and ovarian cancer [38, 39]. In PDAC, down-regulation of miR-497 was associated with poor prognosis [40]. As a tumor suppressor, miR-497 inhibited cell proliferation, gemcitabine sensitivity and apoptosis [41]. Collectively, our findings suggested that miR-497 can be a novel therapeutic target in modulating the HOTTIP level in PDAC. Recently, we developed a high through-put screening method to discover small molecule miRNA modulator [42]. This screening method may be used in developing novel miR-497 modulator, which in turn inhibited HOTTIP in PDAC.

In summary, HOTTIP was ectopically expressed in cancer, including PDAC. HOTTIP promoted PDAC development via canonical *cis*-acting HOTTIP-HOXA13 axis, and non-canonical *trans*-acting HOTTIP-WDR5-MLL1-H3K4met3 pathways (Fig. 6). Under these two pathways, we identified nine novel targets under the regulation of HOTTIP. Novel therapeutic approaches could be developed by interfering the HOTTIP-regulating pathways and its downstream effectors. In addition, restoration of miR-497 that inhibited HOTTIP may be potential therapeutic strategies for PDAC.

**Fig. 6.**
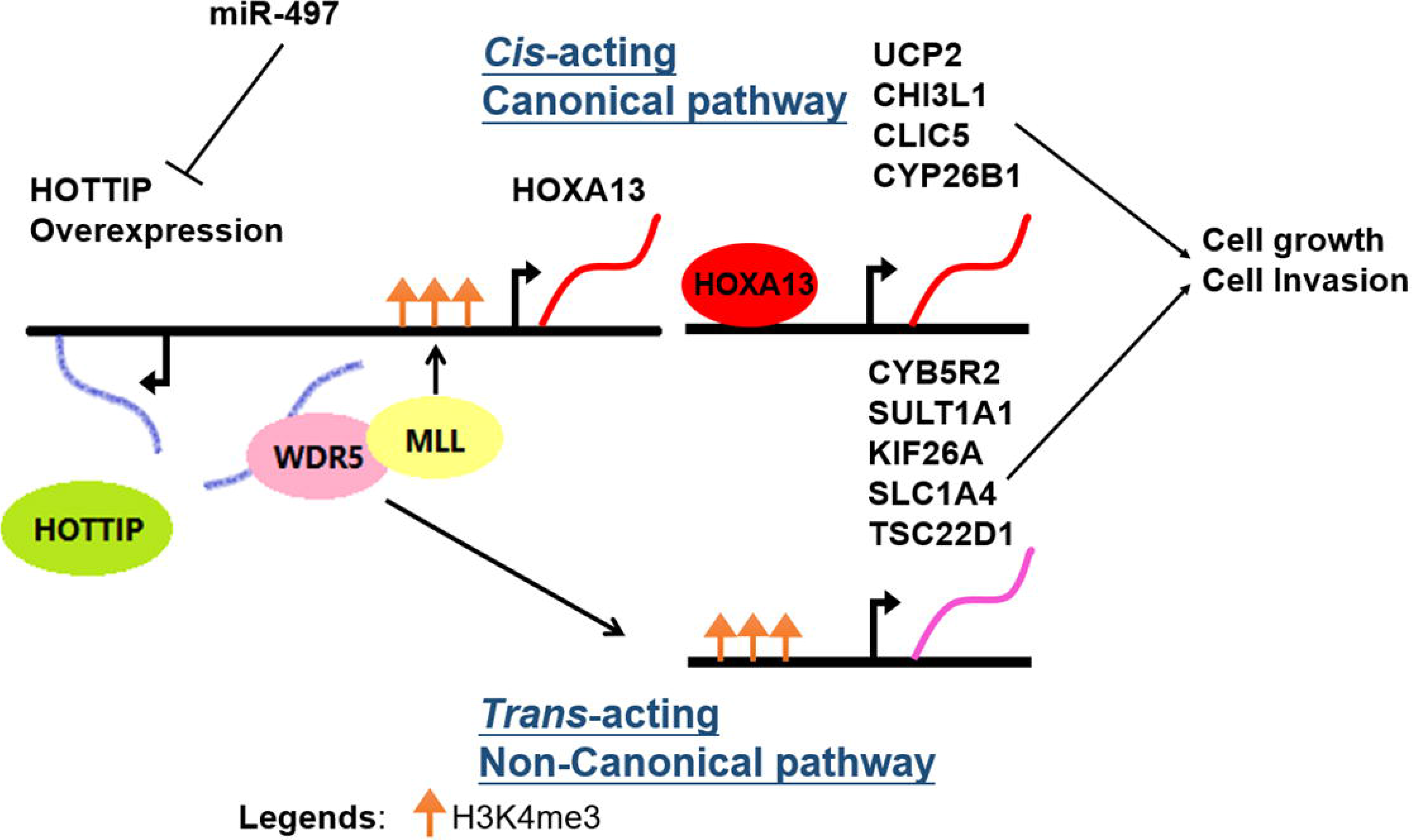
Schematic diagram describing the role of miR497-HOTTIP in the development and progression of PDAC. In PDAC, HOTTIP, which was inhibited by miR-497, was overexpressed. HOTTIP promoted PDAC development via canonical HOTTIP-HOXA13 axis and non-canonical *trans*-acting HOTTIP-WDR5-MLL1 complex. In HOTTIP-HOXA13 axis, HOTTIP activated the expression of HOXA13 which in turn activated CYP26B1, CLIC5, CHI3L1 and UCP2. In HOTTIP-WDR5-MLL1 pathway, HOTTIP-WDR5-MLL1 complex directly activated CYB5R2, SULT1A1, KIF26A, SLC1A4 and TSC22D1 in different chromosomes, without the involvement of HOXA13. These pathways resulted in enhanced cell growth and invasion in PDAC.

## Abbreviations used in this paper

HOTTIP: HOXA transcript at the distal tip
PDAC: pancreatic ductal adenocarcinoma
lncRNA: long non-coding RNA
miRNA: microRNA
miR-497: microRNA-497
CYP26B1: Cytochrome P450 Family 26 Subfamily B Member 1
CLIC5: Chloride Intracellular Channel 5
CHI3L1: Chitinase 3 Like 1
UCP2: Uncoupling Protein 2
CYB5R2: Cytochrome B5 Reductase 2
SULT1A1: Sulfotransferase Family 1A Member 1
KIF26A: Kinesin Family Member 26A
SLC1A4: Solute Carrier Family 1 Member
TSC22D1: TSC22 Domain Family Member 1
WDR5: WD Repeat Domain 5
MLL: Mixed-Lineage Leukemia
H3K4me3: trimethylation of histone H3 at lysine 4
NC: negative control
Ctrl: control

## 5. Acknowledgements

This work was supported by grants from General Research Fund, Research Grants Council of Hong Kong (CUHK462713, 14102714, 4171217 and 14120618 to Y.C.); National Natural Science Foundation of China (81672323 to Y.C.); Direct Grant from CUHK to YC.

## 5.2 Author contribution

CHW and CHL designed the experiments; CHW, CHL and QH generated the data; CHW drafted the manuscript; SLC helped to obtain research materials; JHMT and KFT provided research materials; YC guided the project and revised the manuscript.

## 5.3 Availability of data and materials

Microarray data are available at ArrayExpress E-MTAB-7305.

## 5.4 Ethics approval and consent to participate

The study was carried out with the approval of the Joint CUHK-NTEC Clinical Research Ethics Committee.

