## Supplementary material for "Ectopic HOTTIP Expression Induces Non-canonical Transactivation Pathways to Promote Growth and Invasiveness in Pancreatic Ductal Adenocarcinoma": suppl data

**Supplementary Methods**

**Quantitative reverse-transcription PCR (qRT-PCR)**

Cell lines RNA was extracted by TRIzol Reagent (Invitrogen). Formalin-fixed, paraffin-embedded (FFPE) sample RNA was isolated by miRNeasy FFPE Kit (Qiagen) according to manufacturer’s protocol. Measurement of gene expression level was performed by qRT-PCR. cDNA was synthesized using High-Capacity cDNA Reverse Transcription Kit (Applied Biosystems). Real-time PCR was performed by ABI 7900HT RealTime PCR system using SYBR Green PCR Master Mix (Applied Biosystems). Relative expression of each target mRNA was normalized to GAPDH mRNA level. For measuring the expression level of miRNAs, TaqMan microRNA assays were used according to the manufacturer’s protocol (Applied Biosystems). Real-time PCR measurements were performed by ABI 7900HT RealTime PCR system using miR-497-specific TaqMan primers (Applied Biosystems). The sequences of primers used in this study are shown in Supplementary Table S1.

**Immunohistochemical Staining**

Immunohistochemical staining was performed using human FFPE PDAC tumor samples. The sectioned tissues were deparaffinized and rehydrated by xylene and a series of graded ethanol. Antigen retrieval was performed using PT module (Thermofisher). Then, IHC was performed using Histostain-Plus IHC Kit, HRP, broad spectrum (Life Technologies, Carlsbad, CA). The sections were probed anti-UCP2 antibody (ab97931; abcam). Sections were counter-stained with hematoxylin, were dehydrated with a series of graded ethanol and xylene, and were mounted. A scoring system, based on the percentage of positive cells and staining intensity under the microscope with 100X magnification, was used to quantify the UCP-2 staining. 4 categories (0, 1, 2, and 3) were demoted as 0%, 1-10%, 10-50%, and >50%.

**MTT cell viability assay**

Cell viability was analyzed by 3-(4,5-Dimethylthiazol-2-yl)-2,5-diphenyltetrazolium bromide (MTT) assay. 2,500 cells were seeded for at least triplicate on a 96 well plate and were allowed to grow for several time-points. At each time point, cells were incubated with 0.65 mg/ml MTT diluted in normal culture medium for 2hrs. Cells were then lysed in DMSO and absorbance at 595nm was measured.

**Invasion assay**

Transwell invasion assay was performed by coating matrigel (354234; Corning) on the upper chamber. On the next day, 2.5X10^4^ cells in DMEM without fetal bovine serum (FBS) were placed on the upper chamber. DMEM supplemented with 10% FBS was injected to the lower chamber. After 3 days, the invading cells were fixed by formaldehyde, were permeabilized by methanol, and were stained by crystal violet.

**Chromatin Immunoprecipitation (ChIP)**

ChIP assay was performed using the EZ-Magna ChIP Hi-sens (Millipore) according to manufacturer’s protocol. Cross-linked chromatin was incubated at 4°C overnight with anti-IgG antibody (Millipore), anti-HOXA13 antibody (ab106503; abcam), anti-WDR5 antibody (ab56919; abcam) and anti-MLL1 antibody (#ABE240; Millipore). The precipitated DNA was quantitated by real-time PCR. The data was normalized by IgG antibody and was compared to siNC if necessary.

**Chromatin Isolation by RNA Purification**

Chromatin Isolation by RNA Purification was performed as described by Chu *et al*. [1]. Briefly, cells were fixed by 4% paraformaldehyde. Then the crosslinked chromatin was sonicated and hybridized with five biotin-labeled oligos antisense to HOTTIP at 37°C for 4 hrs with rotation. The hybridized HOTTIP-chromatin was captured by streptavidin-labeled beads, and RNA and DNA were subsequently purified. HOTTIP-binding chromatin was detected by real-time PCR. The data was normalized to GAPDH (negative control).

**Analysis of patient data sets**

Publicly available Pancreatic Ductal Adenocarcinoma patient dataset was obtained from The Cancer Genome Atlas (TCGA). Processed data in log_2_(FPKM-UQ+1) was used in analyzing the expression of HOTTIP and HOXA13. Patients with zero expression of HOTTIP or HOXA13 were excluded. For the association analysis between HOTTIP and HOXA13 expression level, linear regression was used.

1. Chu C, Quinn J, Chang HY. Chromatin isolation by RNA purification (ChIRP). J Vis Exp. 2012, pii: 3912.

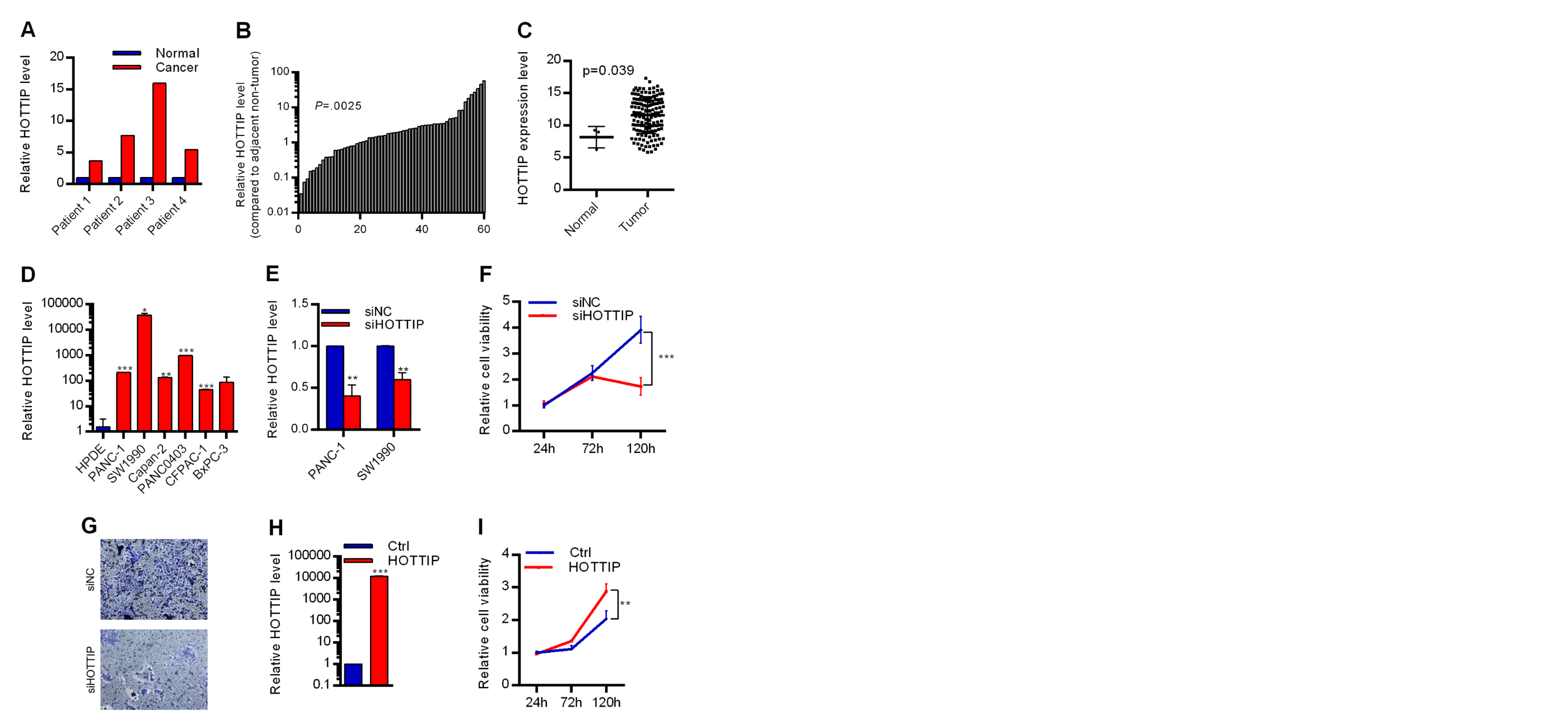

**Fig. S1.** HOTTIP promotes PDAC cell growth *in vitro*. **A**, Genome-wide lncRNA expression microarray was performed to measure the lncRNA expression level in 4 pairs of human PDAC tumor and non-tumor tissues. HOTTIP was upregulated in all PDAC tumor samples. **B**, Expression of HOTTIP in 60 pairs of PDAC tumor samples, compared to adjacent non-tumor tissues. **C**, TCGA analysis of HOTTIP expression in PDAC. **D**, Expression of HOTTIP in the PDAC cell line panel, compared with HPDE cells. Expression was normalized to GAPDH mRNA (n=3). **E**, Knockdown efficiency of siRNA against HOTTIP (siHOTTIP) in PDAC cells. **F**, Knockdown of HOTTIP inhibited cell viability in PANC-1 cells (n=4). **G**, Knockdown of HOTTIP inhibited cell invasiveness in SW1990 cells, demonstrated by transwell cell invasion assay. **H**, Overexpression efficiency of HOTTIP in HPDE cells. **I**, Overexpression of HOTTIP promoted cell growth in HPDE cells (n=4). *P < .05; **P < .01; ***P < .001 when compared with control.

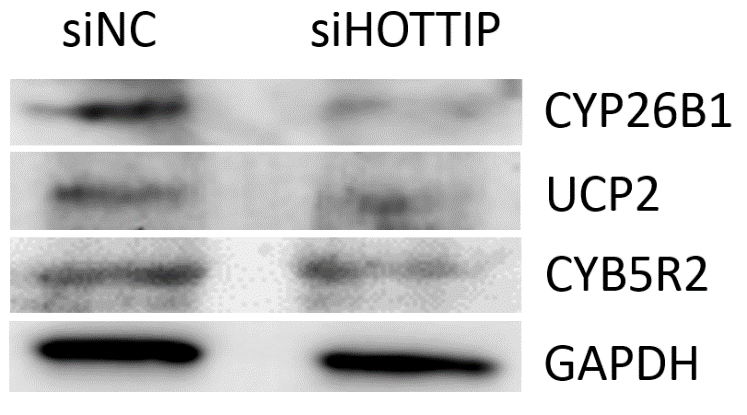

**Fig. S2.** Western blot analysis of CYP26B1, UCP2 and CYB5R2 after knockdown of HOTTIP in PANC-1 cells.

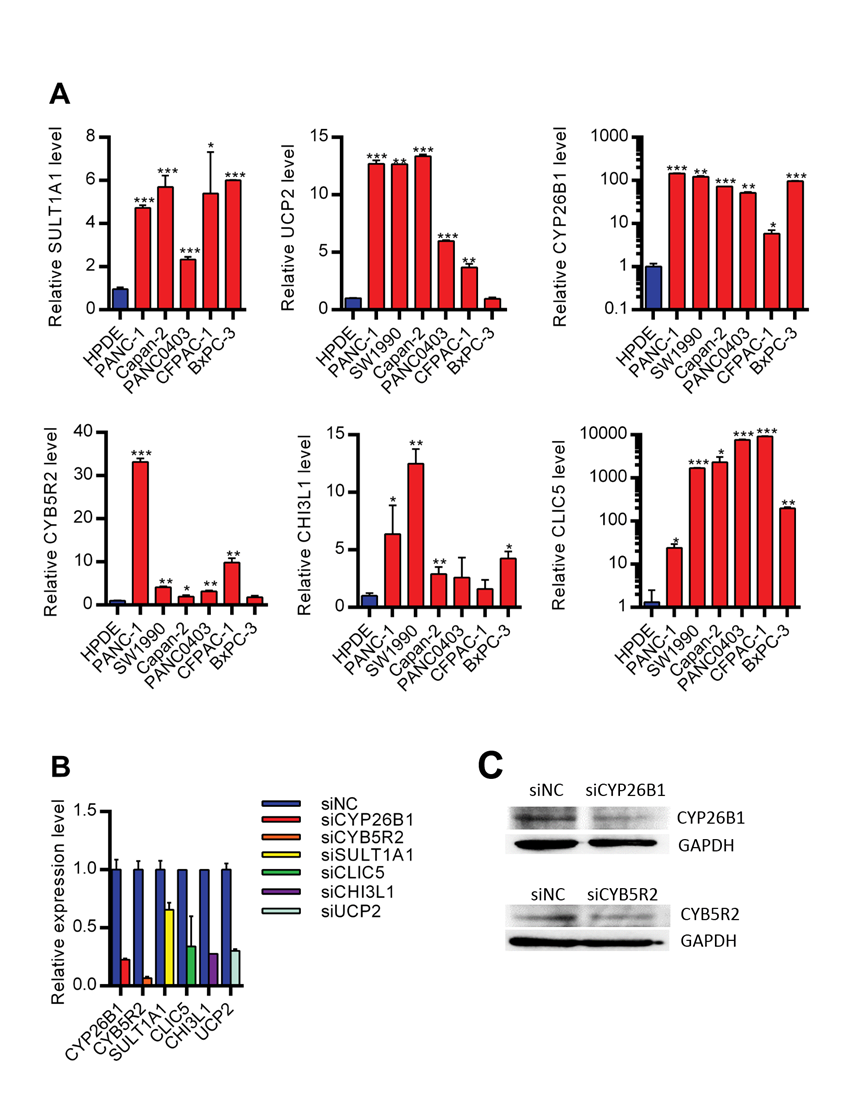

**Fig. S3.** HOTTIP target genes are upregulated in PDAC cells. **A**, Expression of HOTTIP target genes in PDAC cell panel. *P < .05; **P < .01; ***P < .001 when compared with HPDE cell. **B**, Knockdown efficiency of siRNAs targeting CYB5R2, CYP26B1, CLIC5, CHI3L1, UCP2 and SULT1A1. **C**, Western blot analysis of knockdown efficiency of siRNA targeting CYB5R2 and CYP26B1 in PDAC-1 cells.

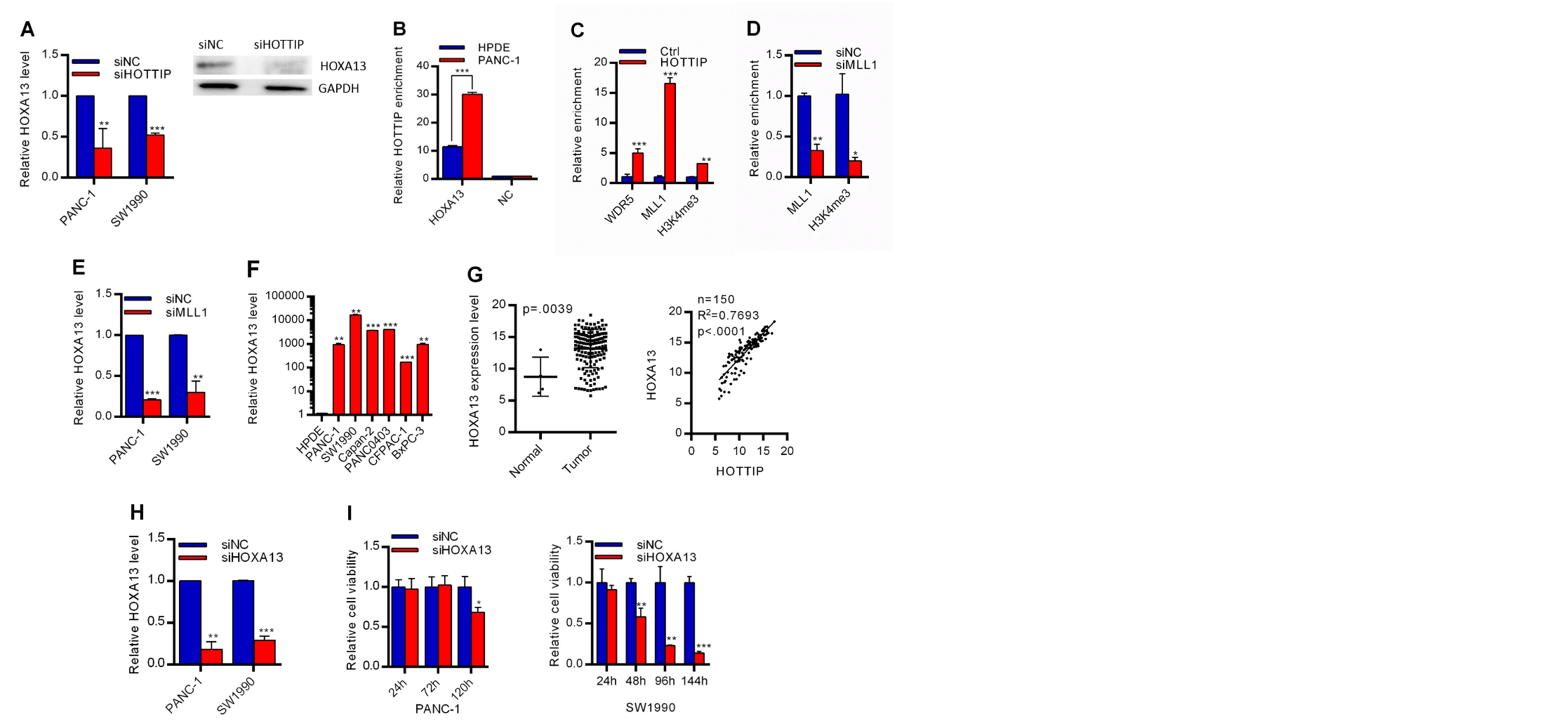

**Fig. S4.** HOXA13 promotes PDAC growth under the regulation of HOTTIP. **A**, HOXA13 expression was reduced in PDAC cells treated with siHOTTIP (n=3). **B**, ChIRP assay analysis of HOTTIP binding to the promoter of HOXA13 in SW1990 cells. Data were normalized to negative control and were compared to HOTTIP enrichment levels in HPDE cells (n=2). **C**, WDR5, MLL1 and H3K4me3 levels at the promoter of HOXA13 were increased when HOTTIP was overexpressed in HPDE cells (n=3). **D**, MLL1 and H3K4me3 occupancies at the promoter of HOXA13 were reduced when SW1990 cells were treated with siMLL1 (n=2). **E**, HOXA13 was down-regulated when HOTTIP was knocked down (n=3). **F**, Expression of HOXA13 in PDAC cell panel, compared to HPDE (n=2). **G**, TCGA analysis of HOXA13 expression and its correlation with HOTTIP expression in PDAC. **H**, Knockdown efficiency of siRNAs targeting HOXA13 (n=3). **I**, Knockdown of HOXA13 inhibited PDAC cell growth (n=4). *P < .05; **P < .01; ***P < .001 when compared with control.

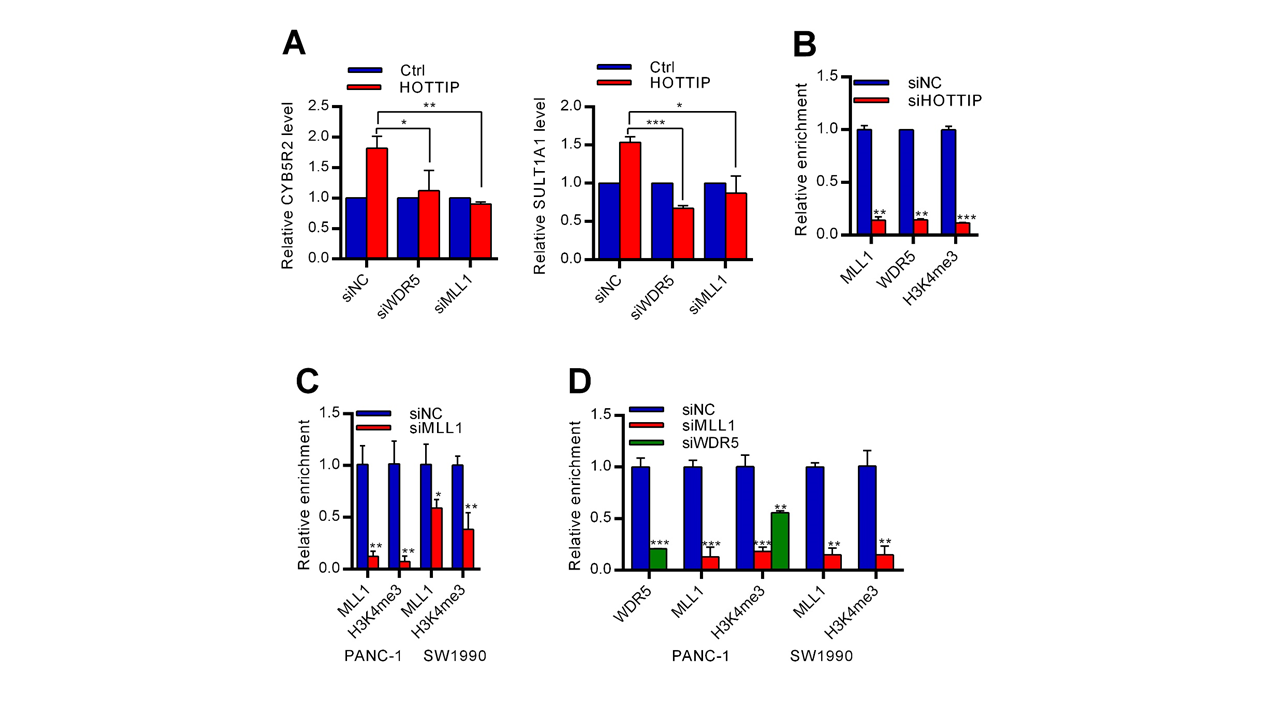

**Fig. S5.** Identification of targets in non-canonical HOTTIP-WDR5-MLL1 pathway. **A**, The increase in expression of CYB5R2 and SULT1A1 by HOTITIP was rescued by knockdown of MLL1 or WDR5 (n=3). **B**, WDR5, MLL1 and H3K4me3 levels at the promoter of CYB5R2 were decreased after knockdown of HOTTIP (n=3). **C**, MLL1 and H3K4me3 levels at the promoter of SULT1A1 in PDAC cells were decreased after knockdown of MLL1 (n=3). **D**, MLL1, WDR5 and H3K4me3 levels at the promoter of CYB5R2 were decreased after knockdown of MLL1 or WDR5 in PDAC cells (n=3). *P < .05; **P < .01; ***P < .001 when compared with control.

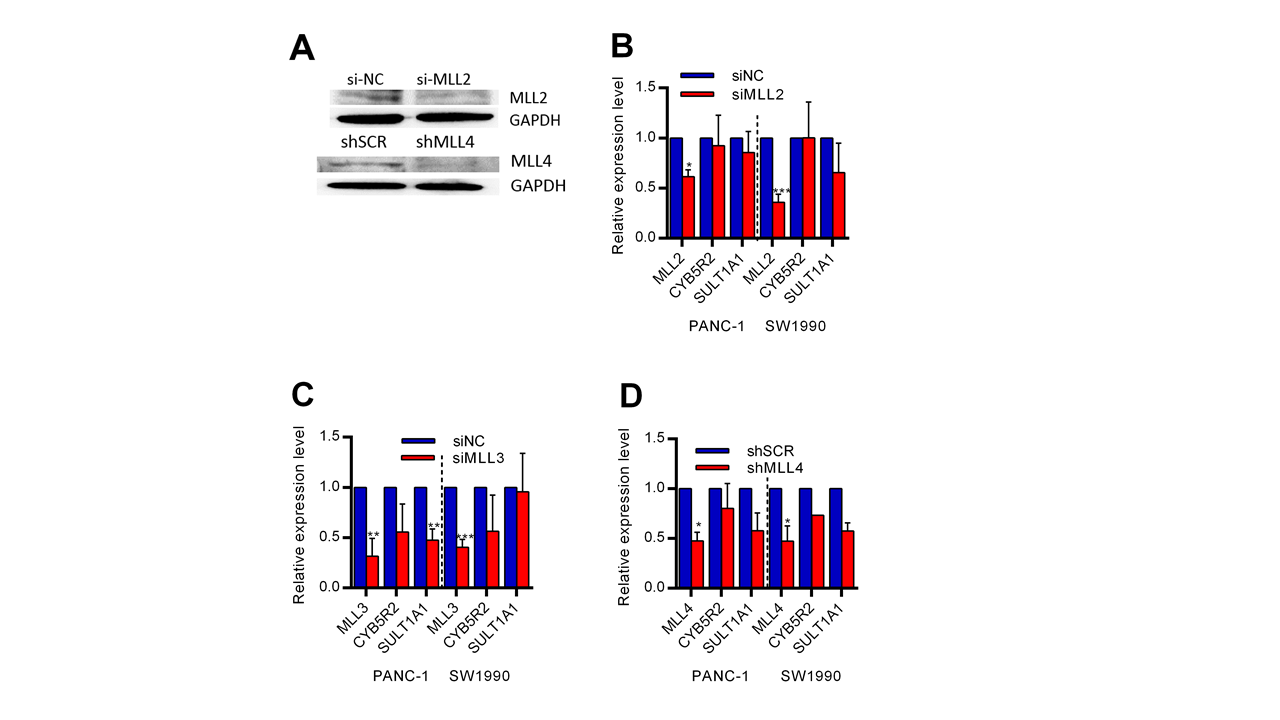

**Fig. S6.** MLL2, MLL3 and MLL4 are not involved in HOTTIP regulatory pathway. **A**, Knockdown efficiency of MLL2 and MLL4. **B-D**, Knockdown of neither (**B**) MLL2, (**C**) MLL3, nor (**D**) MLL4 did not affect the expression of CYB5R2 or SULT1A1 (n=3).

*P < .05; **P < .01; ***P < .001 when compared with control.

**Table S1. Primers used in this study.**

|  | Sequence | Function |
| --- | --- | --- |
| HT-001 qRT-PCR-F | ACGCATATTCACGCATCA | qRT-PCR |
| HT-001 qRT-PCR-R | TTACCAAGCCACAGGAGA | qRT-PCR |
| HOXA13 qRT-PCR F | TGGAACGGCCAAATGTACTG | qRT-PCR |
| HOXA13 qRT-PCR R | TGGCGTATTCCCGTTCAAGT | qRT-PCR |
| CHI3L1-qRTPCR-F | CCACAGTCCATAGAATCCTCGG | qRT-PCR |
| CHI3L1-qRTPCR-R | TGCCTGTCCTTCAGGTACTGCA | qRT-PCR |
| CYP26B1-qRTPCR-F | GCACCTCTTTGAGGTCTACCAG | qRT-PCR |
| CYP26B1-qRTPCR-R | AGGATCTGCCGAGCCTGAATGC | qRT-PCR |
| UCP2-qRTPCR-F | TGGTCGGAGATACCAAAGCACC | qRT-PCR |
| UCP2-qRTPCR-R | GCTCAGCACAGTTGACAATGGC | qRT-PCR |
| CYB5R2-qRTPCR-F | CTCATTCGCCACATCACCAAGG | qRT-PCR |
| CYB5R2-qRTPCR-R | GGTGTACCACAGGTTGAACTGG | qRT-PCR |
| SULT1A1-qRTPCR-F | GGAGTTCATGGACCACAGCATC | qRT-PCR |
| SULT1A1-qRTPCR-R | CCTGCCATCTTCTCCGCATAGT | qRT-PCR |
| CLIC5-qRTPCR-F | TCTGTTGCCCAAGCTCCATGTG | qRT-PCR |
| CLIC5-qRTPCR-R | GCATAGGCGTTCTTGAGGTACC | qRT-PCR |
| KIF26A-qRTPCR -F | AGGTACAGGTTGCCTCTTCC | qRT-PCR |
| KIF26A-qRTPCR -R | CCCCAGAACACCTTCCCTAG | qRT-PCR |
| SLC1A4-qRTPCR -F | TCTTGTGGTTGCAGCTTTCC | qRT-PCR |
| SLC1A4-qRTPCR -F | CCACTCCTAACACCAGAGCA | qRT-PCR |
| TSC22D1-qRTPCR -F | TGCGTGCTCCAAGTACTACA | qRT-PCR |
| TSC22D1-qRTPCR -R | ACACCAGCAGCACTAGGAAT | qRT-PCR |
| MLL1 qRTPCR-F | GTGCTTTGTGGTCAGCGGAAGT | qRT-PCR |
| MLL1 qRTPCR-R | TGTGAGACAGCAACCCACGGTG | qRT-PCR |
| MLL2 qRTPCR-F | GGAATGGGTAGCTCTTTGGCGA | qRT-PCR |
| MLL2 qRTPCR-R | TGCCGAATCAGCAGCTCTCGTA | qRT-PCR |
| MLL3 qRTPCR-F | AGATCAGCGTGGACCCTATCCT | qRT-PCR |
| MLL3 qRTPCR-R | CTCTTGACTCGGCATGGTACCA | qRT-PCR |
| WDR5 qRTPCR-F | AGTGCCTCAAGACTTTGCCAGC | qRT-PCR |
| WDR5 qRTPCR-R | CGATGAGCGTCTTCAGGCACTG | qRT-PCR |
| HOXA13-ChIP-F | TCGCTCGCTCTATCTCAAAGT | ChIP |
| HOXA13-ChIP-R | GAGGCTTGGACTGCATTTAGG | ChIP |
| CYB5R2-ChIP-F | GCAAATTCTTGCCTCAGGAC | ChIP |
| CYB5R2-ChIP-R | GTTTAAGCCGCTCAGTCTGC | ChIP |
| SULT1A1-ChIP-F | ATCCTTTGAGGGCAGGAGTT | ChIP |
| SULT1A1-ChIP-R | CGCACCACCAAGTCTGAGTA | ChIP |
| CHI3L1-ChIP-F | TTTTTGCAATTTACATGCTGA | ChIP |
| CHI3L1-ChIP-R | CGAGCTTGCAAAAGATCCTC | ChIP |
| CLIC5-ChIP-F | CCACCCAAATTAGGGACTCA | ChIP |
| CLIC5-ChIP-R | CGCTTTTGTTGTGCTCAATTC | ChIP |
| CYP26B1-ChIP-F | GACTGGGGTGCAACTTTGTT | ChIP |
| CYP26B1-ChIP-R | CAAACCCACCAGCTTGACTC | ChIP |
| UCP2-ChIP-F | GTAACTGACGCGTGAACAGC | ChIP |
| UCP2-ChIP-F | GTCTTTGGGACTCCGTTTCC | ChIP |
| KIF26A-ChIP-F | AATTAATTCCGGAGCTGAGTTCC | ChIP |
| KIF26A-ChIP-R | CACTTTAGGTCTAACTGTCGCG | ChIP |
| SLC1A4-ChIP-F | CGTGAGAGCTCTGTTTCTTCTAG | ChIP |
| SLC1A4-ChIP-F | ATGGATGAATAAGGGGAGCCTTC | ChIP |
| TSC22D1-ChIP-F | TTTTAAGGTTTGTGGCTCTACGG | ChIP |
| TSC22D1-ChIP-R | AGCGGTACTGGGAGGATTTTATA | ChIP |

**Table S2. siRNAs and miRNA mimics used in this study.**

|  | SENSE 5'-3' | ANTI-SENSE 5'-3' |
| --- | --- | --- |
| HOTTIP | GUACGGAAGUUCCAUUAAUTT | AUUAAUGGAACUUCCGUACTT |
| HOXA13 | GAUAUCAGCCACGACGAAUCUCUCU | AGAGAGAUUCGUCGUGGCUGAUAUC |
| WDR5 | GUGGAAGAGUGACUGCUAATT | UUAGCAGUCACUCUUCCACTT |
| MLL1 | CGAUCAAAUGCCCGCCUAATT | UUAGGCGGGCAUUUGAUCGTT |
| MLL2 | GCCGGAGUUUGUAAUCAAATT | UUUGAUUACAAACUCCGGCTT |
| MLL3 | CCAGCUAUGUACAGAAUUATT | UAAUUCUGUACAUAGCUGGTT |
| UCP2 | CUCCCAAUGUUGCUCGUAATT | UUACGAGCAACAUUGGGAGTT |
| CYP26B1 | CCGUGUUCAAAGACGUAATT | UUCACGUCUUUGAACACGGTT |
| CLIC5 | GUGUCAGUCUAUACCAUUATT | UAAUGGUAUAGACUGACACTT |
| CHI3L1 | GGAGCCACAGUCCAUAGAATT | UUCUAUGGACUGUGGCUCCTT |
| CYB5R2 | GAGGCUUUGUGGACCUAAUTT | AUUAGGUCCACAAAGCCUCTT |
| SULT1A1 | GAGUGUGCGAAUCAAACCUTT | AGGUUUGAUUCGCACACUCTT |
| miR-497 | CAGCAGCACACUGUGGUUUGU | AAACCACAGUGUGCUGCUGUU |

**Table S3. GO Enrichment Analysis after knockdown of HOTTIP in SW1990 cells.**

| Function | Type | Enrichment p-value | % genes in group that are present |
| --- | --- | --- | --- |
| nucleus | cellular component | 5.73E-06 | 1.85409 |
| protein binding | molecular function | 8.96E-06 | 1.82241 |
| zinc ion binding | molecular function | 7.52E-05 | 2.2584 |
| DNA binding | molecular function | 8.71E-05 | 2.33393 |
| metal ion binding | molecular function | 0.000286 | 1.9708 |
| sequence-specific DNA binding transcription factor activity | molecular function | 0.00036 | 2.66963 |
| ubiquitin-dependent protein catabolic process | biological process | 0.000374 | 5.71429 |
| negative regulation of defense response to virus | biological process | 0.000461 | 66.6667 |
| cytoplasm | cellular component | 0.000599 | 1.71067 |
| regulation of cAMP metabolic process | biological process | 0.000914 | 50 |
| regulation of transcription, DNA-dependent | biological process | 0.001462 | 2.41449 |
| polynucleotide adenylyltransferase activity | molecular function | 0.002247 | 33.3333 |
| nucleoplasm | cellular component | 0.002561 | 2.49066 |
| hemopoiesis | biological process | 0.002757 | 8.51064 |
| intracellular | cellular component | 0.003496 | 1.97585 |
| iron ion transport | biological process | 0.004024 | 11.5385 |
| transcription, DNA-dependent | biological process | 0.004562 | 2.7668 |
| cell fate determination | biological process | 0.006523 | 20 |
| response to molecule of bacterial origin | biological process | 0.006523 | 20 |

**Table S4. Top20 dysregulated pathways sensitive to knock-down of HOTTIP in SW1990 cells.**

| Pathway Name | Database | Enrichment p-value | % genes in pathway that are present |
| --- | --- | --- | --- |
| Lysine degradation | kegg | 0.016962 | 6.12245 |
| Non-small cell lung cancer | kegg | 0.021956 | 5.55556 |
| Circadian rhythm - mammal | kegg | 0.024535 | 9.09091 |
| Oocyte meiosis | kegg | 0.035767 | 3.57143 |
| D-Glutamine and D-glutamate metabolism | kegg | 0.043905 | 25 |
| Spliceosome | kegg | 0.057784 | 3.05344 |
| Bladder cancer | kegg | 0.079397 | 4.7619 |
| Synthesis and degradation of ketone bodies | kegg | 0.096124 | 11.1111 |
| Amino sugar and nucleotide sugar metabolism | kegg | 0.099748 | 4.16667 |
| Mineral absorption | kegg | 0.10327 | 4.08163 |
| mTOR signaling pathway | kegg | 0.110417 | 3.92157 |
| Transcriptional misregulation in cancers | kegg | 0.130547 | 2.28571 |
| Sulfur metabolism | kegg | 0.135869 | 7.69231 |
| Shigellosis | kegg | 0.144029 | 3.33333 |
| Terpenoid backbone biosynthesis | kegg | 0.155091 | 6.66667 |
| Cell cycle | kegg | 0.157408 | 2.43902 |
| Glioma | kegg | 0.163534 | 3.07692 |
| Renin-angiotensin system | kegg | 0.173892 | 5.88235 |
| p53 signaling pathway | kegg | 0.175458 | 2.94118 |
| Pancreatic cancer | kegg | 0.183485 | 2.85714 |

.
